# Membrane insertion of chromogranin B for granule maturation in regulated secretion

**DOI:** 10.1101/2019.12.28.890053

**Authors:** Gaya P. Yadav, Haiyuan Wang, Joke Ouwendijk, Mani Annamalai, Stephen Cross, Qiaochu Wang, D. Walker Hagan, Clayton Mathews, Edward A. Phelps, Paul Verkade, Michael X. Zhu, Qiu-Xing Jiang

**Affiliations:** Department of Microbiology and Cell Science, University of Florida, 1355 Museum Drive, Gainesville, FL 32611-0700, USA; Department of Integrative Biology and Pharmacology, McGovern Medical School, The University of Texas Health Science Center at Houston, 6431 Fannin St., Houston, TX 77030, USA; School of Biochemistry, Biomedical Sciences Building, University Walk, University of Bristol, Bristol, BS8 1TD, UK; Department of Pathology, College of Medicine, University of Florida, 1275 Center Drive, Gainesville, FL 32611, USA; Wolfson Bioimaging facility, Biomedical Sciences Building, University Walk, University of Bristol, Bristol, BS8 1TD, UK; J. Crayton Pruitt Family Department of Biomedical Engineering, University of Florida, Gainesville, FL 32611, USA

**Keywords:** Regulated secretion, granule acidification, anion channel, cargo maturation, proinsulin-insulin conversion, insulin-secretory granules, catecholamine loading, dopamine, neurotransmitter release, type 2 diabetes, neurodegenerative diseases, neuroendocrine cancer

## Abstract

Regulated secretion serves responses to specific stimuli in eukaryotes. An anion conductance was found essential for maturation and acidification of secretory granules four decades ago, but its genetic identity was unknown. We now demonstrate that chromogranin B (CHGB), an obligate granule protein, constitutes the long-sought anion channel. High-pressure freezing immuno-electron microscopy and biochemical assays showed native CHGB in close proximity to secretory granule membranes, and its membrane-bound and soluble forms both reconstituted Cl^-^ channels. Release of secretory granules delivered CHGB clusters to plasma membranes, which dominate whole-cell anion conductance. Intragranular pH measurements and cargo maturation assays found that CHGB channels supported proinsulin - insulin conversion and dopamine-loading in neuroendocrine cells. β-cells from *Chgb^-/-^* mice exhibited significant granule deacidification, accounting for hyperproinsulinemia, altered glucose-tolerance response and lower dopamine concentration in chromaffin granules in these animals. Membrane insertion of well-conserved CHGB is thus indispensable for granule maturation in exocrine, endocrine and neuronal cells.

**Highlights:**

- Native CHGB is amphipathic and distributes in the lumen and membranes of secretory granules with contrastingly different destinies and functions.
- Native CHGB, once delivered to cell surface via granule exocytosis, dominates anion conductance in plasma membranes.
- CHGB channels facilitate granule acidification and cargo maturation in cultured and primary neuroendocrine cells.
- CHGB channels from bovine, rat and mouse cells all serve the long-missing, intra-organellar anion shunt pathway in the secretory granules for regulated secretion.

## INTRODUCTION

Regulated secretion is conserved in the eukaryotic kingdom of life. As a major measure to respond to external stimuli, eukaryotic cells rely on regulated secretion to generate local feedbacks or remote regulations within a multicellular system or a commune of unicellular organisms ^1^. In human, exocrine and endocrine cells, neural cells, astrocytes, mesenchymal stem cells, etc. may all utilize regulated secretion to achieve physiological homeostasis and regulation ^2–6^. These cells produce specific bioactive cargo molecules that are densely packed in membrane-enclosed intracellular compartments, whose mature forms are called dense-core secretory granules (DCSGs). DSCGs differ from the lysosome-related organelles (LROs) ^7^. Upon reception of specific secretagogues, DSCGs are exocytosed to release their contents. The released cargo molecules, such as peptide hormones, adrenaline, dopamine, etc. can act locally on cells near the releasing sites (autocrine or paracrine), or those far away through the circulation system (endocrine).

Both positive and negative signals from a variety of cell types are integrated together to achieve a steady state at the systems level. Physiological studies have revealed multiple layers of regulations through regulated secretion, making it critically important for every multicellular tissue or organ inside a human body. At the level of individual cells, regulated secretion is finely tuned. A well-known example is the two phases of insulin release after glucose challenge ^8,9^. But how each endocrine cell achieves a temporally-controlled, quantitative secretion remains largely unresolved.

Biogenesis of nascent secretory granules, sorting and maturation of immature granules, and exocytotic release of mature granules are three main steps of regulated secretion ^1,10^. Granule biogenesis exhibits cell specificity and relies on various molecular players and signals in cells of varying tissue origins ^11–13^; however, its molecular mechanism remains unclear ^14^, despite the appealing hypothesis of protein aggregates-induced membrane budding and possible recruitment of receptors and lipids (*e.g*. cholesterol) at exit sites of the *trans* Golgi network (TGN) ^10,15–20^. Exocytotic release of mature DCSGs utilizes Ca^2+^-triggered membrane fusion catalyzed by specific SNARE complexes ^21–24^, whose fundamental principles are well understood. Normal maturation of secretory granules includes both homotypic fusion of immature secretory granules (ISGs) and budding and removal of missorted or unwanted molecules. This maturation process is accompanied by interior acidification from pH ∼6.5 to ∼5.5 ^16,25^. The molecular mechanisms for sorting or removal of granular contents, luminal acidification, maturation of cargos and packaging / concentrating of granular components are only partially defined.

Prior studies, mainly in chromaffin granules, suggested that normal maturation of nascent ISGs into DCSGs requires a vacuolar-type H^+^-ATPase (vATPase; also called vesicular ATPase) to transport protons and a Cl^-^ conductance to neutralize charge accumulation resulted from proton translocation ^25–28^. The genes for the subunits of the proposed vATPase have been cloned, and its physiological functions are well characterized ^29,30^. The proposed Cl^-^ conductance, however, has remained a mystery for more than 40 years. Organellar cation and anion channels have been identified in all major intracellular membranes ^31–33^. Inositol-1,4,5-trisphosphate receptors (IP_3_Rs), K^+^ channels, and CLC-type Cl^-^ channels / transporters were proposed to reside in secretory granule membranes, but these observations have conflicts or variations due to experimental conditions. Even though earlier studies found no cation permeation in chromaffin granules ^27^, patch clamp recordings from isolated granules revealed different K^+^ channels, IP_3_Rs, a large-conductance Cl^-^ channel and a few CLC-family members ^34–38^. Due to inevitable contamination of isolated secretory granules by other intracellular membranes, the physical locations of these identified channels or transporters in intracellular membranes remain uncertain. In addition, counter-arguments have been raised on whether luminal granular proteins are sufficient to bind and shield all protons and buffer granular pH at an acidic level. Such a mechanism is unlikely because without neutralization, the charged protons bound by cargo molecules cannot be shielded by media or granular proteins, and should lead to a high positive potential (> +60 mV after only a small number of protons cross the membrane) ^25,27^. A shunt pathway is thus a must when large quantities of protons are required to acidify the high contents of biomolecules inside DCSGs. Identifying the unknown shunt conductance with high certainty is therefore critically important and will be the focus of this paper.

Members of the granin superfamily share low similarity in primary structures ^39^ and form distinct phylogenetic subfamilies. Native CHGB aggregates strongly at low pH and with mM Ca^2+ 40,41^. Partially purified CHGB binds strongly to lipid vesicles under certain conditions ^40^. A “tightly membrane-associated form” of CHGB was observed on the surface of PC-12 cells after granule release, which resisted high-salt extraction, but was dissolved with detergents, indicating that some of the rat CHGB protein might be integrated in membranes ^42^. Genome-wide association studies identified multiple mutations in human *Chgb* gene that are closely linked to Type 2 diabetes (T2D; supplementary Table S1) ^43^, heart failure ^44–46^, neurodegenerative diseases and psychiatric disorders ^47,48^. Mechanistically, indirect evidence implicated that intracellular CHGB might form proteinaceous aggregates at the TGN and drive the budding of nascent granular vesicles ^19,49^, although the genuine mechanism is yet to be defined. Studies of *Chgb*–*null* mice by three groups reported tissue-specific variations regarding CHGB’s uncertain physiological roles in granule biogenesis ^12,13,50–52^. The physiological functions of intracellular CHGB in regulated secretion remains largely unknown.

Recently we demonstrated that recombinant murine CHGB (mCHGB) alone suffices to form a highly selective Cl^-^ channel in artificial membranes ^14^. Given the unknown identity of the Cl^-^ conductance in the regulated secretory pathway and the high conservation of CHGB proteins from zebra fish to human, it is natural to ask whether native CHGB may be inserted in granular membranes, function as an anion conductance and support normal granular acidification. Here we investigated the intracellular function of the CHGB channel from six different aspects to elucidate its physiological roles of its membrane-bound form in granule acidification and maturation.

## RESULTS

### CHGB puncta on INS-1 cell surfaces after exocytotic release of secretory granules

The granin proteins in endocrine cells have long been considered and treated as soluble proteins ^1,53^. Although this general view has been challenged by some experimental results ^40,42,54^, it was thought that the insoluble CHGB inside secretory granules resulted from pH-dependent association with membranes and partners. Surprisingly, our recent studies demonstrated that after reconstitution nearly all recombinant mouse CHGB (mCHGB) protein purified from *Sf9* cells are not peripherally associated with membranes, but instead are integrated to form anion channels ^14^. We thus asked whether native CHGB, despite multiple post-translational modifications including glycosylation, phosphorylation and tyrosine-O-sulfation ^55,56^ behaves the same as the recombinant protein.

We first tested if native CHGB is retained on the INS-1 cell surface after the release of secretory granules. INS-1 cells were treated with a high [KCl] solution to release readily releasable granules (RRGs) via activation of voltage-gated Ca^2+^ channels (VGCCs) and Ca^2+^-dependent exocytosis. The cells were then labeled with an anti-CHGB antibody under non-permeabilized conditions followed by an Alexa 488-conjugated secondary antibody. Confocal fluorescence microscopy (FM) at the focal level of the top cell surface revealed multiple rat CHGB (rCHGB) puncta (Fig. 1A), demonstrating that after exocytosis rCHGB remained clustered, presumably, at the exocytotic sites. This resembles the “tightly membrane-associated” rCHGB on PC-12 cells ^42^. From quantification, the high KCl-treated cells have on average ∼19 puncta in the image planes, significantly higher than the control cells (2-3) or those cells stimulated with high KCl but labeled with an antibody against the intracellular suppressor of cytokine signaling 1 (SOCS1) (∼6) (Figs. 1B-C). Similarly, stimulation of INS-1 cells in suspension significantly increased surface labeling by the CHGB antibody (supplementary Figs. S1A, B). A good fraction of native CHGB is therefore retained on the surface of INS-1 cells upon granule release.

**Figure 1.**
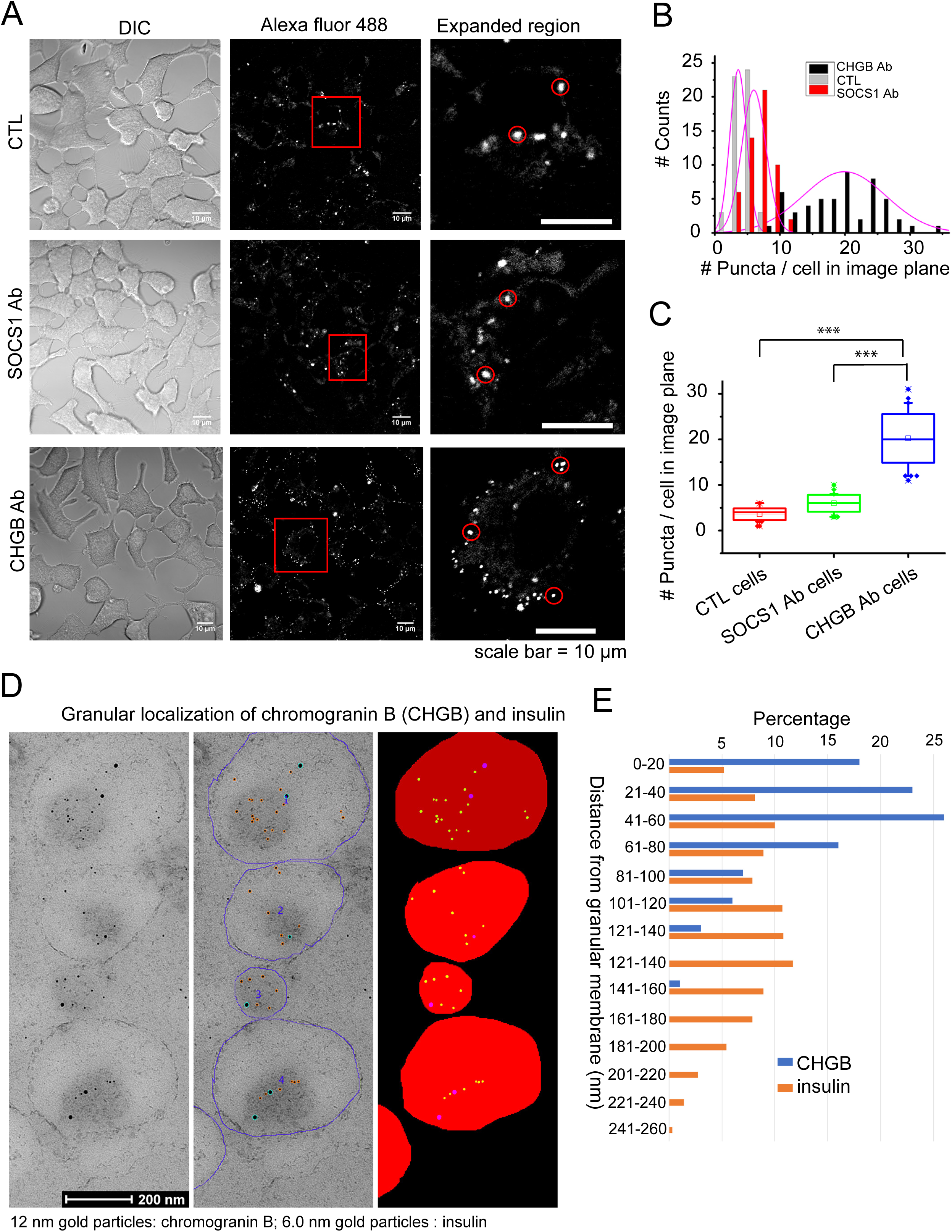
Membrane association of native CHGB on cell surface and inside insulin-secretory granules. (**A**) CHGB retention on cell surface after granule release. INS-1 cells were incubated with high KCl (55 mM) for 15 min at 37 °C to release granules before being labeled with anti-CHGB antibody on ice. An Alexa 488-conjugated secondary (2^nd^) antibody was used to visualize CHGB on cell surface by confocal FM. **Upper row**: images of control (CTL) cells treated only with 2^nd^-Ab; **Middle row**: images of cells labeled with anti-SOCS1 antibody and the 2^nd^-antibody; **Lower row**: images of cells labeled with anti-CHGB antibody and the 2^nd^-antibody. (**B**) Histogram of the number of CHGB puncta per cell within the confocal image planes (top surfaces of the cells) under different conditions (>50 cells) in (A). (**C**) Average number of puncta per cell on the top surfaces. ***: *p* < 0.001 (**D**) HPF-immuno-EM images showing distribution of CHGB and insulin inside DCSGs of rat pancreatic β-cells. CHGB was marked by 12 nm gold particles while insulin by 6 nm ones. Individual granules were demarcated for quantitative analysis (middle and right panels). **(E)** Histogram of CHGB- and insulin-labeling nanogold particles in shortest distances from granular membranes. A good fraction of CHGB-labeling gold particles are in close proximity to the granular membrane as compared to the insulin-labeling ones, which distributes more evenly in DCSGs.

The presence of CHGB puncta on cell surface suggested that other membrane-associated components of the secretory granules might not diffuse away quickly after exocytosis, either. To verify this point, we further demonstrated that vATPase, another major component in the membranes of secretory granules, colocalizes with CHGB on the cell surface as puncta (Fig. S1C). The clustered co-distribution of vATPase and CHGB apparently makes it convenient to retrieve granular membranes and proteins from cell surface via endocytosis ^57^.

Next, we asked whether the native CHGB retained on cell surface became membrane-attached only during or after the granule release, but was fully soluble in the lumen before exocytosis. We performed high-pressure freezing and immuno-electron microscopy (HPF immuno-EM) to compare the distribution of native rCHGB with that of insulin in rat β-cells. HPF immuno-EM is superior in keeping intact original cellular structures and revealing genuinely physiological distribution of target molecules. Dual immunogold labelling of CHGB and insulin was performed on high-pressure-frozen and Lowicryl HM20 freeze-substituted isolated rat islets of Langerhans ^58–60^. CHGB was labeled with 12-nm gold particles and insulin with 6.0-nm ones (left column in Fig. 1D).

For quantitative analysis, the circumference of each granule was manually drawn (middle column in Fig. 1D) before an automated plugin for Fiji was applied to mark the gold particles and fill up the volume of the granule (right column in Fig. 1D). This allowed for automated calculation of the shortest distance from each gold particle to the outer edge of the granule within the section plane. A total of 324 granules from three independent experiments were analyzed (Fig. 1E). As no obvious difference was observed between individual experiments, all data from three were pooled. The average granule diameter was 296 ± 82 nm (mean ± SD), which is slightly larger than previously observed ^60^. The distribution plots of nearest distances to granular membranes, grouped in 20-nm bins, were obviously different between CHGB and insulin (Fig. 1E). While insulin, soluble in granular lumen, showed an relatively even distribution with an average distance of 116.6 ± 62.2 nm (mean ± SD), CHGB exhibited an obviously skewed distribution towards membrane, with ∼70% 12-nm gold particles situated at ≤ 60 nm from the nearest granular membranes and an average distance of 52.1 ± 32.4 nm (p<0.00001, *t*-test with unequal variances). Given the dimensions of the 1^st^ and 2^nd^ antibodies (∼25-30 nm together), the length of conjugation linkers and the sizes of the nanogolds, CHGB molecules integrated in the membrane should be within ∼50 nm from the granular membranes (Fig. 1E). These results indicate that contrary to insulin, native CHGB is more preferentially present at or near the membranes of insulin-secretory granules in rat pancreatic β-cells. The native CHGB thus probably exists in both membrane-associated and soluble forms inside the granules.

### Direct recordings of granular CHGB channels retained on cell surface

To reconcile the membrane association of CHGB with its well-assumed solubility, Pimplikar and Huttner proposed that CHGB must interact with other partners in membranes ^42^. We recently showed that when reconstituted in artificial membranes *in vitro*, recombinant mCHGB alone suffices to form a chloride channel ^14^. This brings up the intriguing possibilities that native CHGB may be inserted into granular membranes and form anion channels, too. The native CHGB puncta in plasma membranes may reflect at least a good portion of membrane-bound proteins, which would become exposed to the extracellular side after soluble granular contents diffuse away. If so, the putative CHGB channels in the puncta should be accessible for electrophysiological recording. To test this prediction, we performed whole-cell patch-clamp recordings of PC-12 cells with pipette and bath [Cl^-^] set at 6.0 mM and 152 mM, respectively. With the cell being held at -80 mV, a voltage protocol made of a ramp from -80 to +80 mV in 100 ms, a 50-ms step at 80 mV and a 400-ms step to 0 mV was applied every second. The depolarization voltages activated VGCCs to facilitate Ca^2+^ influx, and thus triggered release of RRGs via Ca^2+^-dependent membrane fusion. Accordingly, we detected an increase of outwardly rectifying currents in the presence of 2.5 mM Ca^2+^ outside, but not when Ca^2+^ was absent (Figs. 2A, 2C). The currents exhibited a negative reversal potential (*b* in Fig. 2B) and the outward currents were largely abolished by removal of extracellular Cl^-^ (Figs. 2A-B), suggesting that they carried mostly Cl^-^. After ∼5 minutes of repetitive stimulation, the current reached a maximum (time point *b* in Fig. 2A), which presumably reflects the full depletion of all RRGs.

**Figure 2.**
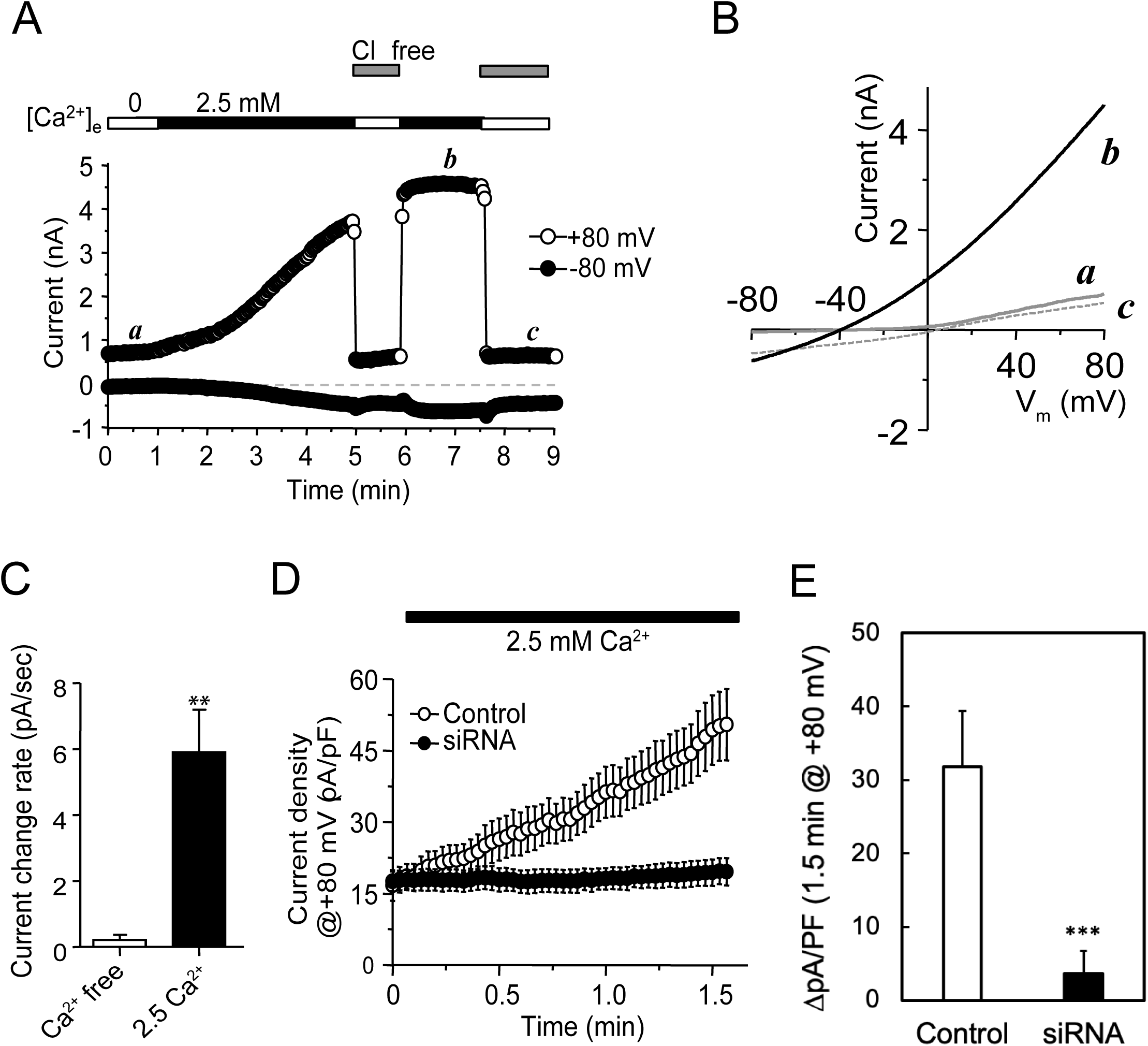
Ion currents through native CHGB channels retained in plasma membranes after granule release in PC-12 cells. (**A**) Whole-cell recordings from a PC-12 cell (top bars; out of more than 10 cells) that was initially perfused with a Ca^2+^-free solution and then with a normal bath containing 2.5 mM Ca^2+^. Pipette electrode contained low Cl^-^. Holding potential was -80 mV. Every second, a voltage protocol made of a ramp (−80 to +80 mV in 100 ms), a step (50 ms at 80 mV) and another step (400-ms at 0 mV) was delivered. Whole-cell currents were stable in Ca^2+^-free bath (first period including time point *a*) and increased gradually in the normal bath. Switching the bath to a Cl^-^-free solution abolished the increased outward current. After 5 minutes, the outward current (+80 mV) became saturated (fourth period including time point *b*), which was almost completely abolished after removal of extracellular Cl^-^ (fifth period including time point *c*). The inward current at -80 mV showed a much smaller Ca^2+^-dependent increase and was only slightly sensitive to extracellular Cl^-^-free solution. (**B**) Typical current traces elicited by ramp pulses at time points *a*, *b*, and *c*. The trace at time point *b* showed a reversal potential of ∼ - 40 mV and weak outward rectification. Traces at time points *a* and *c* had a reversal potential of ∼ 0 mV and almost no rectification. (**C**) Average current change rate (mean ± SEM) at +80 mV, expressed as pA/sec, for cells in Ca^2+^-free (n = 5) and 2.5 mM Ca^2+^ (n = 14) solutions. **: *p* < 0.05 by Student’s *t* test. (**D**) Time course of current development at +80 mV for cells transfected with scrambled (control; open circles) and CHGB-targeting siRNAs (siRNA; solid circles). Pipette solution contained NMDG-Cl. Cells were held at 0 mV. Voltage ramps from -80 to +80 mV in 500 ms with a 50-ms step at -80 mV before and a 50-ms step at +80 ms after the ramp, were applied every 2 sec. Cells were placed in Ca^2+^-free bath first and then in a normal bath with 2.5 mM Ca^2+^. Data points are mean ± SEM of current densities at +80 mV from 6 Control- and 7 CHGB siRNA-transfected cells. (E) Comparison of changes in average current density over 1.5 minutes for the same groups of cells showed in (**D**). ***: p < 0.005.

The residual currents after substitution of extracellular Cl^-^ with gluconate could stem from either some cation conductance or some, albeit weak, permeation of gluconate through the anion channels. Either of the two would cause a right shift of the experimental reversal potential from the calculated Nernst potential of ∼-82 mV for Cl^-^. From the measured reversal potential of -40 mV, Cl^-^ conductance would be 5 times of that for monovalent cations (including leak) if there was no gluconate permeation, or the anion channel had a Cl^-^: gluconate permeability ratio of 6:1 if all recorded currents were from the same type of anion channels. Realistically, in the presence of small leak and gluconate permeation, Cl^-^ currents should contribute to more than 85% of the measured current increase after RRG release (Fig. 1A).

To assess if granular CHGB is responsible for the exocytotic increase in Cl^-^ conductance in the plasma membrane, we knocked down CHGB expression before making electrical recordings (Fig. 2D; supplementary Figs. S2A-C) ^12^. CHGB siRNAs suppressed CHGB expression by > 98% (Figs. S2A-B). Among four CHGB siRNAs in the mixture, each alone efficiently knocked down CHGB, with siRNAs 2 and 4 being ∼2-fold more effective than the other two (Fig. S2C). Thus, siRNAs 2 and 4 were used either individually or in combinations in different experimental repeats. The same results from these repeats suggested no obvious OFF-target effects. We hence pooled together data from cells treated by individual and combined siRNAs in quantitative analysis. In CHGB knockdown INS-1 cells, we also compared the release of granules after treatment with high [KCl] by detecting vATPase on cell surfaces (Fig. S2D vs. Fig. 1A), and found that similar to pancreatic hormone release in islets isolated from *Chgb^-/-^* mice ^13^, CHGB knockdown still allows delivery of vATPase in surface puncta, though to a lesser extent, after depolarization-triggered granule release.

Next we compared the development of depolarization-induced Cl^-^ currents in PC-12 cells transfected with control (sequence scrambled) or CHGB siRNAs. This time, an NMDG-Cl pipette solution was employed to minimize outward cation currents. While the currents increased steadily over time in cells transfected with the control siRNAs, they never did so in those transfected with the CHGB siRNAs (Fig. 2D-E). These data demonstrate that the appearance of Cl^-^ conductance on the cell surface following the release of RRGs in PC-12 cells is dependent on the expression of CHGB protein.

Notably, CHGB knockdown significantly decreased the number of intracellular granules in INS-1 (Figs. S3A, S3E) and PC-12 cells that were stained with DND-160 (Fig. S3F). In PC-12 cells, a ∼50% decrease in the average number of secretory granules per cell after CHGB knockdown is comparable to the ∼30% reduction in granule content released from *Chgb-null* chromaffin cells ^61^. Because granule exocytosis remained normal in *Chgb-/-* cells (Fig. S2D) ^50,52^, a 50% decrease in granule number was expected to incur ∼50% loss of Cl^-^ currents in the CHGB-knockdown cells should the Cl^-^ conductance be contributed by another protein. However, we observed nearly 100% loss of Cl^-^ currents, congruent with CHGB directly contributing to the Cl^-^ conductance carried to plasma membranes by RRGs. If the single channel conductance of a native CHGB channel (in 150 mM Cl^-^) is similar to that of the recombinant one, ∼125 pS ^14^, at least 300 channels ought to be delivered to produce a steady-state, whole-cell Cl^-^ current of ∼4.5 nA at +80 mV.

### Both membrane-bound and soluble CHGB proteins from bovine pancreatic secretory granules are capable of reconstituting anion channels

Although native CHGB functions as a Cl^-^ channel on cell surface after granule release (Fig. 2) and recombinant CHGB forms a Cl^-^ channel *in vitro* ^14^, it remained unresolved whether CHGB in fully sealed secretory granules inside a cell indeed functions as a channel, instead of forming a channel during or after exocytic membrane fusion, and if so, whether it serves as the long-sought anion conductance that supports granule acidification. In native cells CHGB undergoes various posttranslational modifications, experiences contrastingly different lipid environments in secretory granules and interacts with various partners and granular cargos, all of which might disrupt its channel function. Because native secretory granules are on average ∼300 nm in diameter (Fig. 1D), too small to be assessed by direct patch-clamp recording, we approached this critical question from three orthogonal aspects: 1) biochemical analysis of native proteins from isolated secretory granules, 2) granule maturation in cultured neuroendocrine cells and 3) granular acidification in pancreatic β-cells of acutely isolated islets of Langerhans.

To evaluate whether native bovine CHGB (bCHGB) inside secretory granules has the capacity to form Cl^-^ channels, we isolated secretory granules from bovine pancreas ^62,63^. The soluble contents inside the granules were released in three freeze-thaw cycles. Roughly equal amounts of bCHGB proteins were detected in soluble and membrane-bound fractions (Fig. S4), in accord with previous reports ^42,53^. In SDS-PAGE, mature bCHGB ran at ∼100 kDa (Figs. S4, 3A-B, 3H-I), probably due to posttranslational modification. In size-exclusion chromatography (SEC), the membrane-bound bCHGB was eluted at the same position as the recombinant mCHGB dimers ^14^. The bCHGB hence behaves biochemically similar to the recombinant protein from *Sf9* cells.

To examine if the purified bCHGB can form Cl^-^ channels, both soluble and membrane-bound forms were purified in detergents and reconstituted into KCl-loaded vesicles to measure channel activity by a light-scattering-based flux assay ^14^. The assay detects increased Stokes radii of vesicles after sudden KCl efflux. In vesicles containing DOPC, sphingomyelin and cholesterol, the membrane-bound bCHGB yielded robust Cl^-^ flux (Fig. 3D, 0.0 mM Cl^-^). Increasing extravesicular [Cl^-^] from 0.2 to 2.0 mM (Fig. 3D) gradually decreased Cl^-^ flux to nil, revealing an apparent Cl^-^-binding affinity of 0.47 mM (Figs. 3D, E), which is almost the same as that of the recombinant mCHGB channels in egg PC vesicles (0.49 mM) ^14^. DIDS, an anion channel blocker, inhibited bCHGB channel activity with an apparent *k_D_* ∼ 0.43 μM and a Hill coefficient, *n,* of ∼1.0 (Figs. 3F-G), also nearly identical to what’s measured for mCHGB. The native bCHGB channel is thus pharmacologically equivalent to the recombinant one.

**Figure 3.**
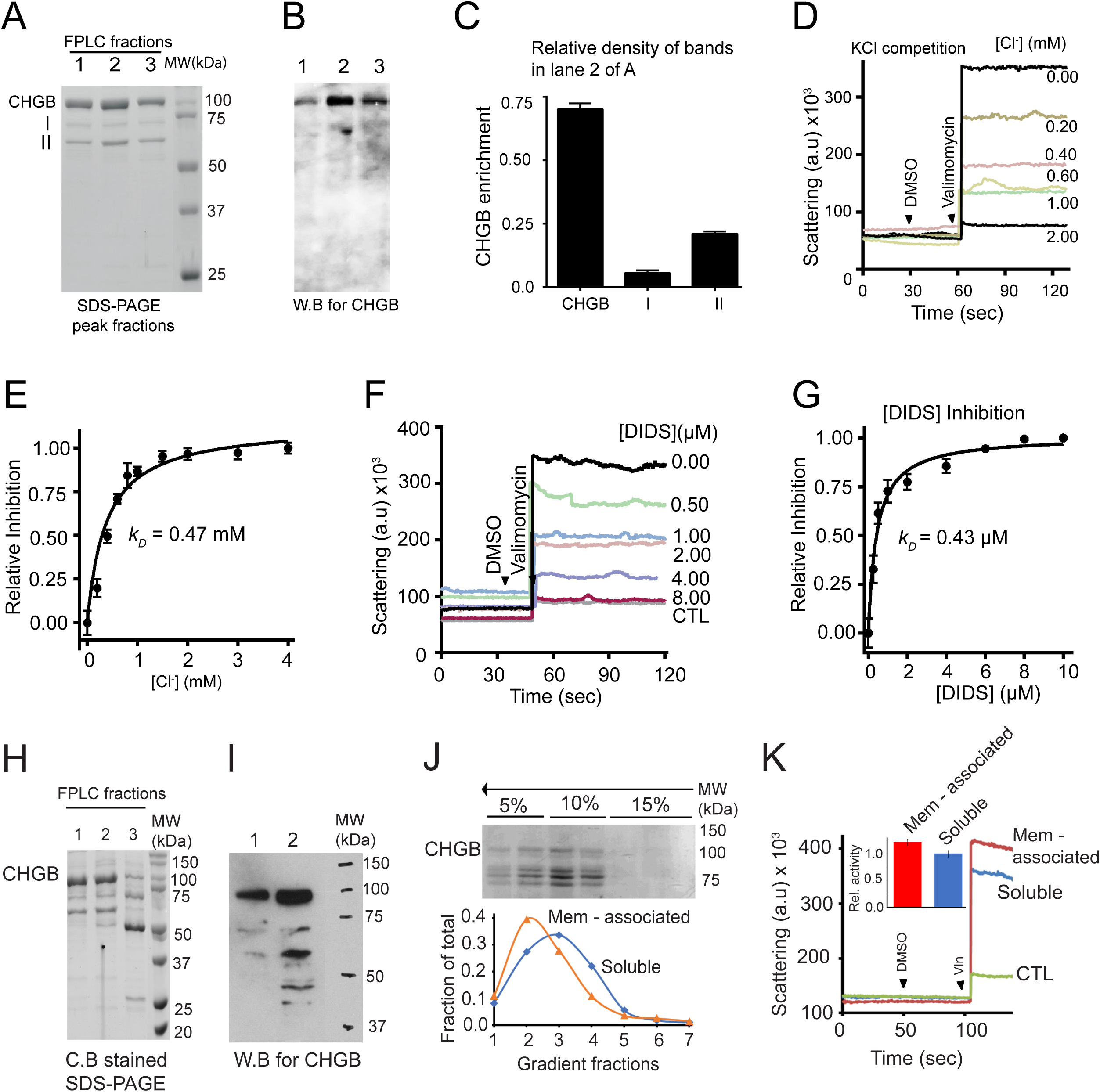
Membrane-bound and soluble bovine CHGB proteins both reconstitute Cl^-^ channels in membrane. (**A**) Purified bCHGB in detergents from membrane-bound fractions of bovine pancreatic granules. bCHGB peak fractions eluted out of a Superdex 200 were assayed by SDS-PAGE and Coomassie blue staining. **(B)** Western-blot of bCHGB from the three fractions in (**A**). (**C**) Quantification of bCHGB and two contaminating bands in lane 2 of (**A**). (**D**) Light-scattering-based flux assay of purified membrane-bound bCHGB reconstituted in vesicles loaded with 300 mM KCl. Vesicles were first exchanged into an external solution containing 300 mM KI and varying amount of KCl. Valinomycin-triggered change in light-scattering was recorded. DMSO was used as control. (**E**) Inhibition of Cl^-^ efflux by extravesicular Cl^-^. Data from (**D**) (black circles; error bars are *s.d.,* n=3) fitted with a Hill-equation (black line) yielded an apparent *k_D_* of ∼0.47 mM and a Hill coefficient of 1.0. (**F&G**) Inhibition of Cl^-^ efflux by DIDS. Fitting of the flux data points from (**F**) (black circles; error bars: *s.d.* n=3) with a Hill-equation yielded a *k_D_* of ∼ 0.43 μM and a Hill coefficient of ∼1.0. **(H)** bCHGB in soluble fractions released from pancreatic granules were eluted from a Superose S6 column. Its three peak fractions were assayed by Coomassie blue-stained SDS-PAGE. The first two fractions contain similar bands as those in (**A**). The third fraction has a clear degradation band at ∼50 kDa. (**I**) Western blot of bCHGB separated as in (**H**). CHGB proteins in soluble fractions exhibited more degradation than those in membrane-associated fractions (**B)**. (**J**) ***<u>Top:</u>*** bCHGB purified from soluble fractions reconstituted in vesicles made of DOPC: sphingomyelin: cholesterol. Vesicles were floated from bottom to top in a Ficoll 400 density gradient (5%,10% and 15%). Gradient fractions were assayed by SDS-PAGE and Coomassie blue staining. Full-length bCHGB is marked. More degradation was observed after reconstitution. ***<u>Bottom</u>***: Full-length bCHGB bands in individual lanes of the top gel were quantified and normalized against total full-length bCHGB in all fractions, and plotted to show the relative distribution of CHGB among different fractions (blue) For comparison, similar analysis of bCHGB purified from membrane-bound fractions in (**A**) were conducted and plotted in orange. Soluble fractions showed more degradation, and was more poorly reconstituted than membrane-bound ones. (**K**) bCHGB from soluble (blue) and membrane-associated (red) fractions were compared in the light-scattering-based assay for Cl^-^ flux. The control vesicles had no protein and showed no signal (green). **Inset:** Relative activity in the ion-flux assays from the two different forms of native bCHGB was compared quantitatively.

As recombinant mCHGB does, the amphipathic nature of bCHGB allows its soluble form to reconstitute Cl^-^ channels in membrane. After purification the soluble bCHGB showed similar band patterns as the membrane-bound form (Fig. 3H vs. 3A), despite more degradation products (lane 2 in Figs. 3H-I). However, the full-length bCHGB still accounts for the majority of the total protein (>80%; Fig. 3I). Reconstitution of soluble bCHGB was incomplete when vesicles were assayed after floatation in a density gradient (top in Fig. 3J) ^14^. Quantification of bCHGB bands in Coomassie blue-stained gels showed slightly higher reconstitution efficiency of membrane-bound CHGB than the protein from the soluble fractions (Fig. 3J, bottom panel, red *vs*. blue traces). Nonetheless, the soluble bCHGB reconstituted in vesicles yielded robust Cl^-^-flux, comparable to that generated by vesicles made with the same amounts of membrane-associated protein (mem-associated vs. soluble; Fig. 3K). Control experiments using empty vesicles yielded negative results. In summary, both soluble and membrane-bound bCHGB proteins extracted from bovine pancreatic granules are equally capable of reconstituting Cl^-^ channels that are pharmacologically equivalent to the recombinant mCHGB channels.

### CHGB channel supports normal granular maturation in insulin-secretory cells

In the second set of experiments, we assessed the possible physiological roles of native CHGB inside secretory granules in cultured cells. We reasoned that if CHGB serves as the long-sought shunt pathway for Cl^-^, disrupting its function should slow down or even abolish acidification and maturation of secretory granules. To test this prediction, we knocked down CHGB expression and measured intragranular pH. INS-1 cells were selected as the first model system because they recapitulate insulin secretion as native pancreatic β-cells and their intragranular pH levels have been extensively studied with a well-established procedure using DND-160, a ratiometric pH sensor ^64^.

We first confirmed that DND-160 stains well the secretory granules in INS-1 cells. A fluorescently-labeled granular protein, syncollin-pHluorin ^65^, was transiently expressed and imaged together with DND-160-stained compartments in live cells (Fig. S4A). By confocal microscopy, DND-160-stained compartments all showed strong signals of syncollin, suggesting that they are indeed secretory granules ^64^. Because of the possible variations in DND-160 loading among different granules, ratiometric measurements from dozens to hundreds of granules (Figs. 4A-D, S3B-D) were acquired in order to compare the distribution of pH values systematically. Furthermore, when INS-1 cells loaded with DND-160 for 5 minutes were compared with those loaded for 30 minutes, the measured distribution of intragranular pH was the same. Dye loading for 5 minutes was thus sufficient for our experiments.

**Figure 4.**
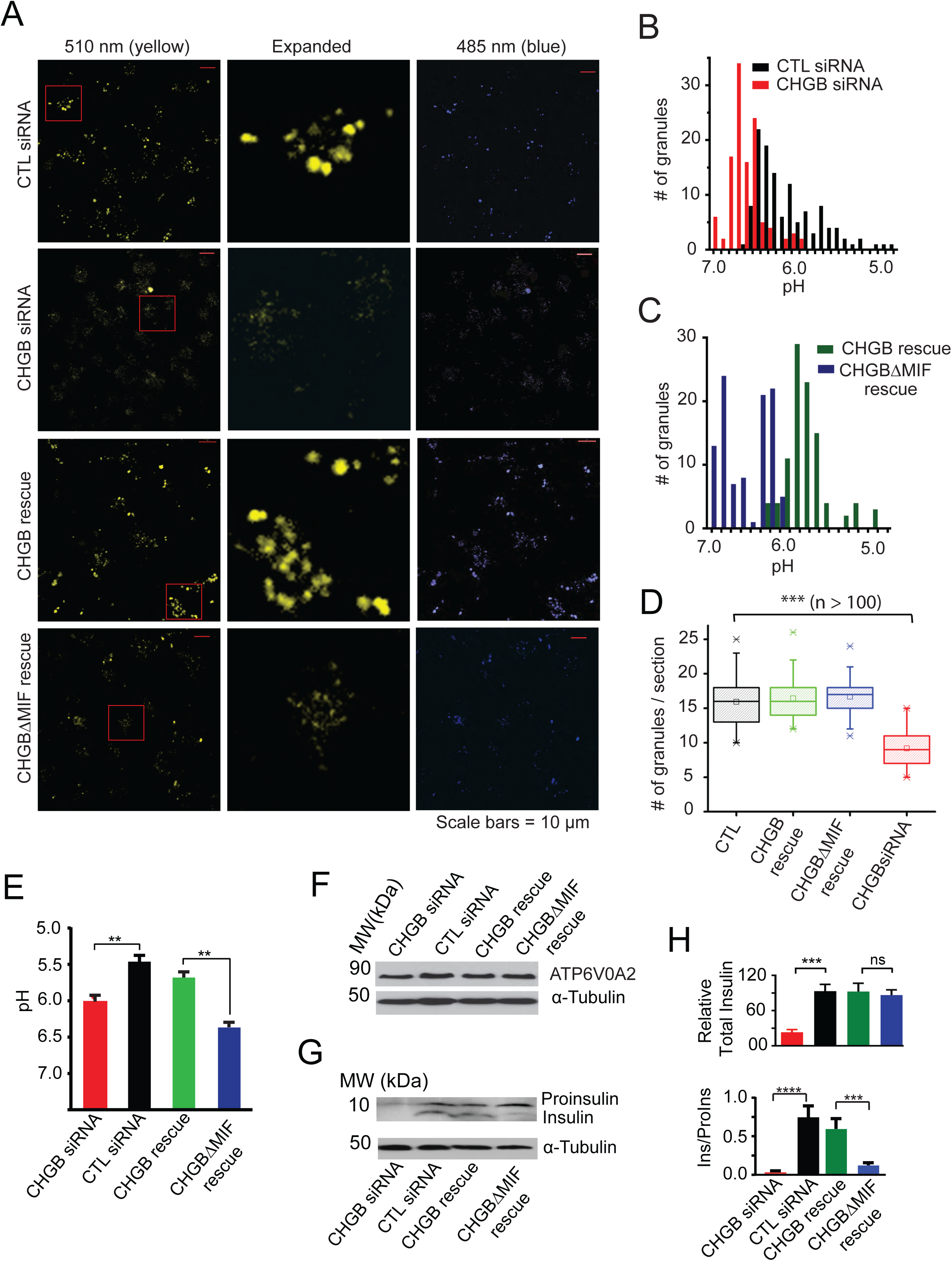
Native CHGB channel is critical for normal granule acidification and insulin maturation in INS-1 cells. (**A**) Ratiometric pH measurements. INS-1 cells in a glass-bottomed dish (DIC) were transfected with control (CTL; top row) or CHGB siRNAs (rows 2 to 4). In top two rows, cells were stained after 96 hours with one medium change at 48th hour. In rows 3 & 4, 48 hours after transfection with CHGB siRNAs, cells were transfected to express CHGB or CHGBΔMIF and were imaged after 48 more hours. Excitation: 410 nm; emission: 485 nm & 510 nm. A small area (red window) in each 510 nm image was expanded to show granules. Measurement at 510-nm in row 4 was weaker due to pH difference. (**B**) Histogram of intragranular pH values from individual cells treated with CTL or CHGB siRNAs (black vs. red). (**C**) Histogram of intragranular pH values from cells whose endogenous CHGB were suppressed by siRNA before transient overexpression of wild-type CHGB or CHGBΔMIF (green vs. blue bars). (**D**) Average number of granules per INS-1 cell in the confocal image plane under different conditions in **(A)**. (**E**) Average pH values of secretory granules (n > 110) from cells as in **(A-C)**. (**F**) Western blot of ATP6V0A2 (vesicular H^+^-ATPases) from cells treated as in **(A**). (**G**) Western blot of insulin and proinsulin from lysates of cells treated as in (**A**). (**H**) Relative total insulin (insulin + proinsulin; upper panel) and insulin maturation (Ins / ProIns; lower panel) from cells treated as in (**A**). Relative total insulin was normalized to cells treated with CTL siRNAs (black). Error bars are *s.d*. in **(D)** (n > 100) and **(G)** (n=3). **: *p* < 0.05; ***: *p* < 0.001; ****: *p* < 0.0001; *ns*: not significant.

To evaluate the importance of CHGB channels to granular acidification, we knocked down CHGB expression in INS-1 cells before overexpressing full-length mCHGB or its non- conducting deletion mutant (CHGBΔMIF). For DND-160, dual-excitation (Figs. S3B-D) and dual-emission experiments (Figs. 4A-D) produced similar results. Comparison of cells transfected with control- (CTL) or CHGB-specific siRNAs showed that CHGB knockdown reduced the total number of secretory granules per cell (Figs. 4A, 4D, S3E-F) and increased average intragranular pH (called deacidification; Figs. 4A-B, S3C-D). In dual excitation experiments, cells treated with NH_4_Cl served as a positive control because NH_4_^+^ increases intragranular pH to a neutral or slightly basic level. The pH distributions among 100-140 granules randomly selected from paired images of multiple cells (details in Supplementary Information) showed an obvious deacidification after NH_4_Cl treatment (Figs. S3C-D), and the CHGB knockdown shifted the pH distribution in the same direction (Figs. 4B, S3C). The average pH in the secretory granules that remained after CHGB knockdown in INS-1 cells (∼30%; Fig. S3E) was significantly higher than that from CTL siRNA-transfected cells (red vs. grey in Fig. S3D). Similar results were obtained from dual-emission experiments (Figs. 4A-B). Therefore, we focused on the dual emission experiments to avoid near-UV excitation.

After transient expression of wild-type CHGB and CHGBΔMIF in CHGB-knockdown cells, we repeated intragranular pH measurement (rows 3 & 4 in Fig. 4A; Figs. 4C-D). Both constructs contained the N-terminal sorting signal (Cys-loop) needed for protein sorting and granule biogenesis ^20^. The overexpressed proteins thus restored the number of granules (rows 3 & 4 vs. row 2 in Fig. 4A, Fig. 4D), confirming that the recombinant mRNAs overwhelmed siRNAs for protein expression. However, ratiometric pH measurements found that only the wild-type CHGB, but not the non-conducting CHGBΔMIF mutant, restored granular acidification (Figs. 4C, E). The overexpressed CHGBΔMIF even showed dominant negative effects (Fig. 4E). These data demonstrate that the channel function of native CHGB, not its activity in granule biogenesis, is required for normal granule acidification in INS-1 cells.

Because vATPase pumps protons, we tested whether CHGB knockdown caused deacidification by reducing its expression. Western blotting showed that ATP6V0A2 expression was not changed by knockdown or by overexpressing CHGB or CHGBΔMIF (Fig. 4F), ruling out the possibility that the impaired granule acidification resulted from decreased vATPase level. As a control, we measured intragranular pH using NPY-ClopHensor ^66^, which was co-expressed with CHGB or CHGBΔMIF for ratiometric measurements (Fig. S5). An *in sit*u calibration curve (Fig. S5B) was obtained to read out intragranular pH values in INS-1 cells overexpressing CHGB and CHGBΔMIF, which on average were ∼5.5 and ∼6.6 respectively (Fig. S5C), similar to the results from DND-160-stained granules (Fig. 4E). As observed in CHGB knockdown cells overexpressing CHGBΔMIF (Figs. 4A, 4E), CHGBΔMIF exhibited a dominant negative effect, likely through oligomerization with endogenous CHGB in forming tetrameric channels and making them nonfunctional. A second control is on granule biogenesis induced by CHGBΔMIF. When overexpressed in HEK293T and *Npc1*^-/-^ CHO cells, both CHGB and CHGBΔMIF and their fusion proteins with ClopHensor induced granule-like vesicles that were stained with DND-160 (Figs. S6A-B), reassuring that CHGBΔMIF is as good as wild-type CHGB in supporting granule biogenesis (Fig. 4A).

Because of lower prohormone convertase activity at elevated pH levels ^67^, loss of granule acidification should impair proinsulin-to-insulin conversion (insulin maturation). Western blotting showed that in contrast to CTL, CHGB knockdown in INS-1 cells drastically reduced total insulin (proinsulin plus insulin) level, which was restored by overexpressing either CHGB or CHGBΔMIF in the knockdown cells (Figs. 4G, 4H). Thus, the effect on total insulin is related to the capacity of CHGB (or CHGBΔMIF) in inducing granule biogenesis (Fig. 4H top). More interestingly, CHGB knockdown impaired insulin maturation, as shown by the lack of mature insulin band (Fig. 4G) and the insulin / proinsulin (Ins / ProIns) ratio being close to nil (Fig. 4H, bottom). While overexpression of wild-type CHGB corrected the defect in insulin maturation, CHGBΔMIF expression did not (green vs. blue; Fig. 4H bottom). Because CHGBΔMIF conducts no Cl^-^, it could not restore granule acidification or insulin maturation. The roles of CHGB in granule acidification and maturation are thus tightly coupled, and match with the characteristics of the unknown Cl^-^ conductance first proposed in chromaffin granules ^25,27^.

### Native CHGB channel facilitates normal dopamine loading in PC-12 cells

To evaluate how generally the CHGB channel functions for granule acidification and maturation, we performed similar experiments in PC-12 cells and compared intragranular pH’s among four differentially treated cells (Figs. S7, 5A-C). Similar to INS-1 cells, CHGB knockdown in PC-12 cells decreased the number of secretory granules (Figs. S3F and S7), which was restored by CHGB or CHGBΔMIF (Fig. S7). CHGB knockdown de-acidified secretory granules in PC-12 cells (Figs. 5A-B), which was reversed by overexpressing wild-type CHGB, but not CHGBΔMIF (Figs. 5A, 5C). Manipulations in PC-12 did not alter ATP6V0A2 expression, either (Figs. 5D). Therefore, like INS-1 cells, PC-12 cells rely on CHGB channel activity for normal granule acidification.

**Figure 5.**
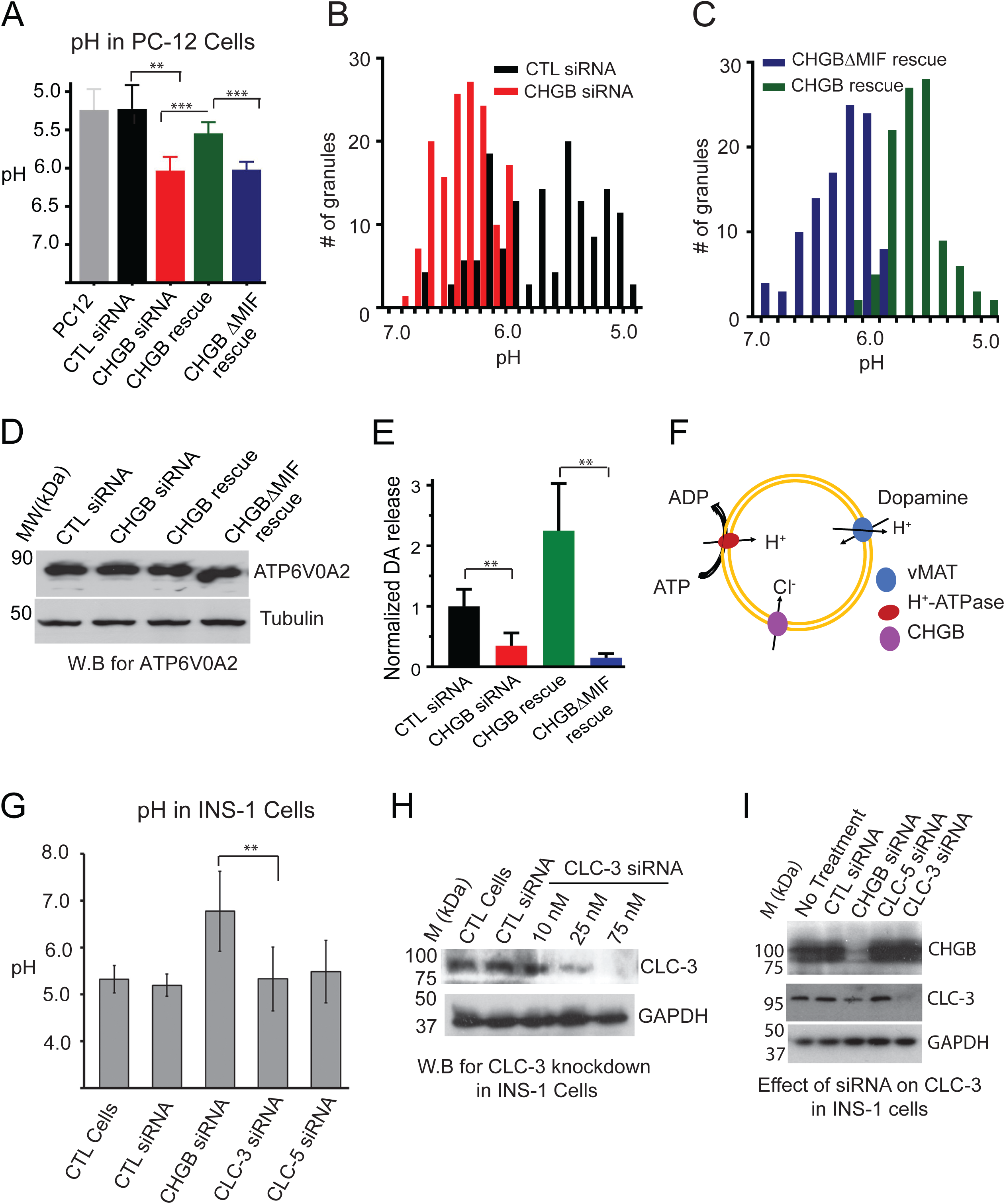
CHGB channel function is critical for granule acidification and dopamine loading in PC-12 cells. (**A**) Ratiometric pH measurements. PC-12 cells were transfected with control (black) or CHGB siRNAs (red); after 48 hours some of the CHGB knockdown cells were transfected to overexpress CHGB (greed) or CHGBΔMIF (purple). After 96 hours, cells were treated with DND-160 for ratiometric imaging. Typical images are showed in supplementary figure S7. Average granular pH values from hundreds of granules are shown in a bar graph. Average intragranular pH from granules in PC12 cells without siRNA treatment is in grey. (**B)** Histogram of intragranular pH of PC-12 cells transfected with control (black), and CHGB (red) siRNAs. (**C**) Histogram of intragranular pH values of CHGB-knockdown PC-12 cells overexpressing wild-type CHGB (green) or CHGBΔMIF (blue). (**D**) Western blot of ATP6V0A2 in four differentially treated PC-12 cells (**B&C**). The same number of cells were used for analysis. (**E**) Relative dopamine content in RRGs released from depolarization-treated cells. An ELISA kit was used. Standard *t*-test for (**A**) and two-tailed Welch’s *t*-test for (**E**). **: *p* < 0.05; ***: *p* < 0.001. (**F**) Schematic drawing to show H^+^ pumping and Cl^-^ influx that drive granular acidification and H^+^-coupled transport of dopamine (or epinephrine) into secretory granules by vMAT. (**G**) CLC-3 or CLC-5 knockdown has negligible effect on intragranular pH. INS-1 cells were transfected with control, CHGB, CLC-3 or CLC-5 siRNA. After 96 hours, cells were loaded with DND-160 for imaging. Average pH values were calculated from hundreds of stained granules, and were presented as a bar diagram. **: *p* < 0.05. There was no significant change in average intragranular pH for cells treated with CLC-3 or CLC-5 siRNAs. (**H**) Western blot shows effective knockdown of CLC-3 in INS-1 cells using different concentrations of siRNAs. (**I**) Western blot showing no obvious cross-effect between the three different siRNAs.

Dopamine is the main cargo inside the secretory granules of PC-12 cells. It is pumped into granules by a H^+^-coupled vesicular monoamine transporter (vMAT) (Fig. 5F) ^68^. Granular deacidification in PC-12 cells is expected to decrease dopamine content in RRGs. ELISA measurements ^69^ showed that CHGB knockdown significantly decreased dopamine content in RRGs, which was recovered by overexpressing wild-type CHGB, but not the non-conducting CHGBΔMIF (Fig. 5E). In accord with our data, decreased granular dopamine content was observed in *Chgb^-/-^* chromaffin cells ^50,61^, although these groups attributed it to abated dopamine-binding capacity of granular matrix. Based on our analysis, granule deacidification-induced suppression of dopamine loading would probably be the real cause.

### CHGB does not function indirectly by altering CLC-3 or -5 expression

Although CLC-3 was known to be absent in the secretory granules, we still examined whether in our experimental systems, the physiological effects of CHGB had contributions from CLC-3 or CLC-5. Among INS-1 cells transfected with siRNAs targeting CHGB, CLC-3 or CLC-5, only CHGB siRNAs caused granule deacidification (Fig. 5G), suggesting that neither ClC-3 nor CLC-5 serves as the key anion channel for granule acidification. By western blotting, the CLC-3 siRNAs were founded highly effective and specific in knocking down CLC-3 expression without affecting CHGB (Fig. 5H vs. 5I). Inversely, CHGB siRNAs had no effect on CLC-3 expression, either (Fig. 5I). Therefore, the effect of CHGB knockdown on granule acidification did not occur by indirectly altering CLC-3 (or -5) expression, but did so by directly suppressing CHGB channels.

### Primary pancreatic β-cells from *Chgb^-/-^* mice display impaired granule acidification

Previously, CHGB-knockout mice were reported to exhibit reduced storage and release of catecholamines from chromaffin cells ^12,13,50^ and a decreased level of stimulated secretion of insulin and other pancreatic hormones from islets (Obermuller et al., 2010). Because pancreatic islets of *Chgb-null* animals appeared unable to compensate well for the CHGB loss (Obermuller et al., 2010), we used them to evaluate whether CHGB knockout leads to impaired granule acidification in primary β-cells, which in turn elicits neuroendocrine phenotypes in the mutant mice. A *Chgb*^-/-^ mouse strain produced by the Wellcome Trust Sanger Institute and distributed by EMMA (Figs. 6 & S8; #10088 at URL: https://www.infrafrontier.eu/) was raised by the UF animal care service. All animal work followed an IACUC-approved protocol. Genotyping was performed by PCR using primers recommended by the Sanger Institute (Fig. S8). Pancreatic islets were isolated from mice of ∼8 weeks or older. Male and female littermates were analysed separately in order to identify any sex-related differences. Intragranular pH in pancreatic β cells was measured either in isolated islets (Figs. 6B-D) or in β-cell monolayers derived from these islets (Figs. 6E-G). Fig. 6A shows representative images of isolated islets. Western blotting of CHGB in liver tissues from *Chgb*^+/+^ and *Chgb*^-/-^ mice confirmed the loss of CHGB protein in the latter. Western blotting was also performed for other tissues to confirm complete penetrance of the knockout.

**Figure 6.**
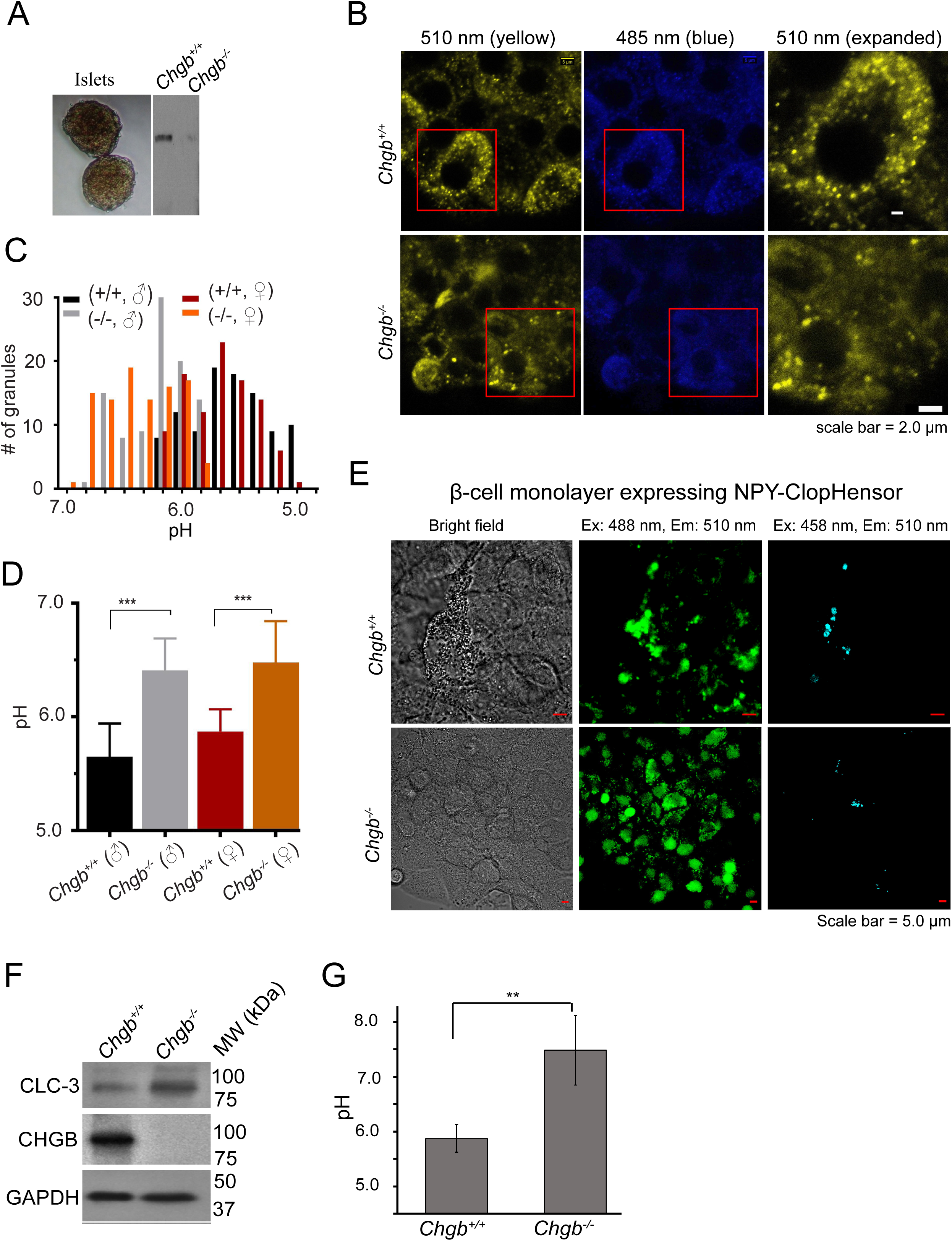
Murine CHGB supports normal granule acidification of insulin-secretory granules in primary pancreatic β-cells. (**A**) Two typical pancreatic islets isolated from mice (left) and western blotting of CHGB from liver tissues removed from a wild-type (*Chgb^+/+^*) and a CHGB-knockout (*Chgb^-/-^*) mouse. (**B**) Typical ratiometric images of pancreatic β-cells in individual islets from *Chgb*^+/+^ and *Chgb^-/-^* mice. Ten islets of each genotype were stained with DND-160 and imaged under a Zeiss LSM-800 confocal microscope (63x objective). Excitation: 410 nm; emission: 485 nm and 510 nm. A small area (red window) in each 510 nm image was expanded to show granules (yellow channel in the right column). β-cells from *Chgb^-/-^* mice have fewer bright granules (above background) and fainter signals at 510 nm due to deacidification. (**C**) Histograms of intragranular pH values measured from granules in isolated islets from both male and female mice of different genotypes. All mice were littermates and were 9-weeks old at the time of analysis. (**D**) Average pH values for four different groups of mice in (**C**). **: *p* < 0.05; ***: *p* < 0.001. (**E**) Typical ratiometric images in monolayers of β-cells grown out of pancreatic islets isolated from *Chgb*^+/+^ and *Chgb^-/-^* mice. After ∼7 days, monolayers of cells were transfected with an NPY-ClopHensor-expressing plasmid, and after another 48-72 hours, were imaged under a Zeiss LSM-710 confocal microscope (63x oil objective). Excitation: 488 nm (green) and 458 nm (cyan); emission: 510 nm. β-cells from *Chgb*^-/-^ mice have fewer granules than the wild-type cells. (**F**) Western blotting of CLC-3 and CHGB in pancreatic tissues from *Chgb*^+/+^ and *Chgb*^-/-^ mice. GAPDH was used as loading control. (**G**) Average intragranular pH measured from ClopHensor expressed in monolayer β-cells from *Chgb*^+/+^ and *Chgb*^-/-^ mice. **: *p* < 0.05.

For whole islet imaging, freshly isolated islets were cultured for two days before DND-160 staining and imaging. To avoid variations in DND-160 staining of cells away from islet surfaces, we focused on cells closer to the surface. Compared to *Chgb*^+/+^ β-cells (top row in Fig. 6B), *Chgb*^-/-^ β-cells contained fewer well-stained, bright granules (bottom row; Fig. 6B), agreeing with decreased granule biogenesis and deacidification as observed in INS-1 and PC-12 cells (Figs. S3E-F, 4E, 5A). The distribution of intragranular pH measured from hundreds of granules showed clear shift to higher values in *Chgb*^-/-^ β-cells from both male and female mice (Fig. 6C). The average intragranular pH values in *Chgb*^-/-^ β-cells are significantly higher by ∼0.7 pH units than those in wild-type cells (Fig. 6D), demonstrating that the lack of CHGB rendered substantial deacidification of insulin-secretory granules in pancreatic β-cells of both sexes (Figs. 6C-6D). Deacidification by ∼0.7 pH units may slow down proinsulin-insulin conversion enough to account for the previously observed increase of proinsulin release from *Chgb*^-/-^ islets ^13^. As a control, we confirmed that in *Chgb*^-/-^ pancreatic islets, CLC-3 expression was not decreased, but instead increased slightly (Fig. 6F). Therefore, the observed granule deacidification in *Chgb^-/-^* β-cells was not secondary to a loss of CLC-3.

As an alternative of pH measurement in islet cells, we prepared cultured β-cell monolayers following an established protocol ^70^ and transiently expressed NPY-ClopHensor fusion protein (Fig. S5) in them for pH measurement (Figs. 6E, 6G). Significant granular deacidification was observed with the difference in average intragranular pH (∼1.5 pH units) between *Chgb*^-/-^ and wild-type cells being greater than that (∼0.7 pH units) from DND-160-stained islet cells (Fig. 6G vs. 6D). These results reassured impaired acidification of secretory granules in *Chgb^-/-^* cells, which underlies the defective insulin maturation in and elevated proinsulin release from the islets of *Chgb^-/-^* mice^12,13,50^.

## DISCUSSIONS

### CHGB-membrane insertion and channel activity in native secretory granules

From six different angles, our data congruently demonstrate a major physiological function of native full-length CHGB in serving as the long-sought Cl^-^ conductance in secretory granules that supports normal granule acidification and maturation. The identity of such a conductance has been elusive for more than four decades and the candidate approach in the past failed to identify it ^25^. The amphipathic nature of the CHGB channel, differing from all known canonical anion channels, made it evade all previous efforts. As far as we know, the channel function is the first well-grounded intracellular function of the CHGB subfamily of granins. Among all amphipathic proteins that form ion channels, CHGB is the first that shows exquisite anion selectivity.

Despite multiple posttranslational modifications, native CHGB remains amphipathic and exists in soluble and membrane-associated forms, both of which reconstitute anion channels in membranes. The native channels retain the same high sensitivity to DIDS and luminal [Cl^-^] (Figs. 3E, 3G). After granule release, the membrane-associated CHGB channels remain as clusters in the plasma membranes and conduct Cl^-^. Because normal cells have sufficient surface area in ER membranes, nascent CHGB could be membrane-associated in ER as well; but it is unclear whether it functions as a channel in ER (or *Golgi*). In DCSGs, CHGB is concentrated and may induce the formation of nanoparticles or nanotubules as the recombinant protein does ^14^. Its soluble forms are likely stabilized by self-oligomerization or forming complexes with other proteins or lipids (Figs. 1D & 3H). The “tightly membrane-associated” CHGB channels keep granular transmembrane potential close to the Nernst potential of Cl^-^ and neutralize luminal positive charges translocated by vATPase. After RRG release, hundreds of CHGB channels are delivered to the surface of each PC-12 cell, making the cell membrane highly permeable to Cl^-^. We postulate that the high Cl^-^ selectivity of the CHGB channel prevents intracellular anionic metabolites from being passively concentrated into secretory granules and being dumped as waste via exocytosis, or when delivered to cell surface, it stops small organic metabolites inside the cell from diffusing out. In *Chgb*^-/-^ cells, the lack of Cl^-^ influx impedes the neutralization of luminal positive charge, stops continued proton pumping ^71^, impairs granule acidification, delays proinsulin-insulin conversion in INS-1 and β-cells (Fig. 7A) and decreases H^+^-coupled pumping of dopamine into secretory granules in PC-12 cells (Fig. 7B). These deficits were corrected by overexpressing wild-type CHGB, but not the non-conducting CHGBΔMIF.

**Figure 7.**
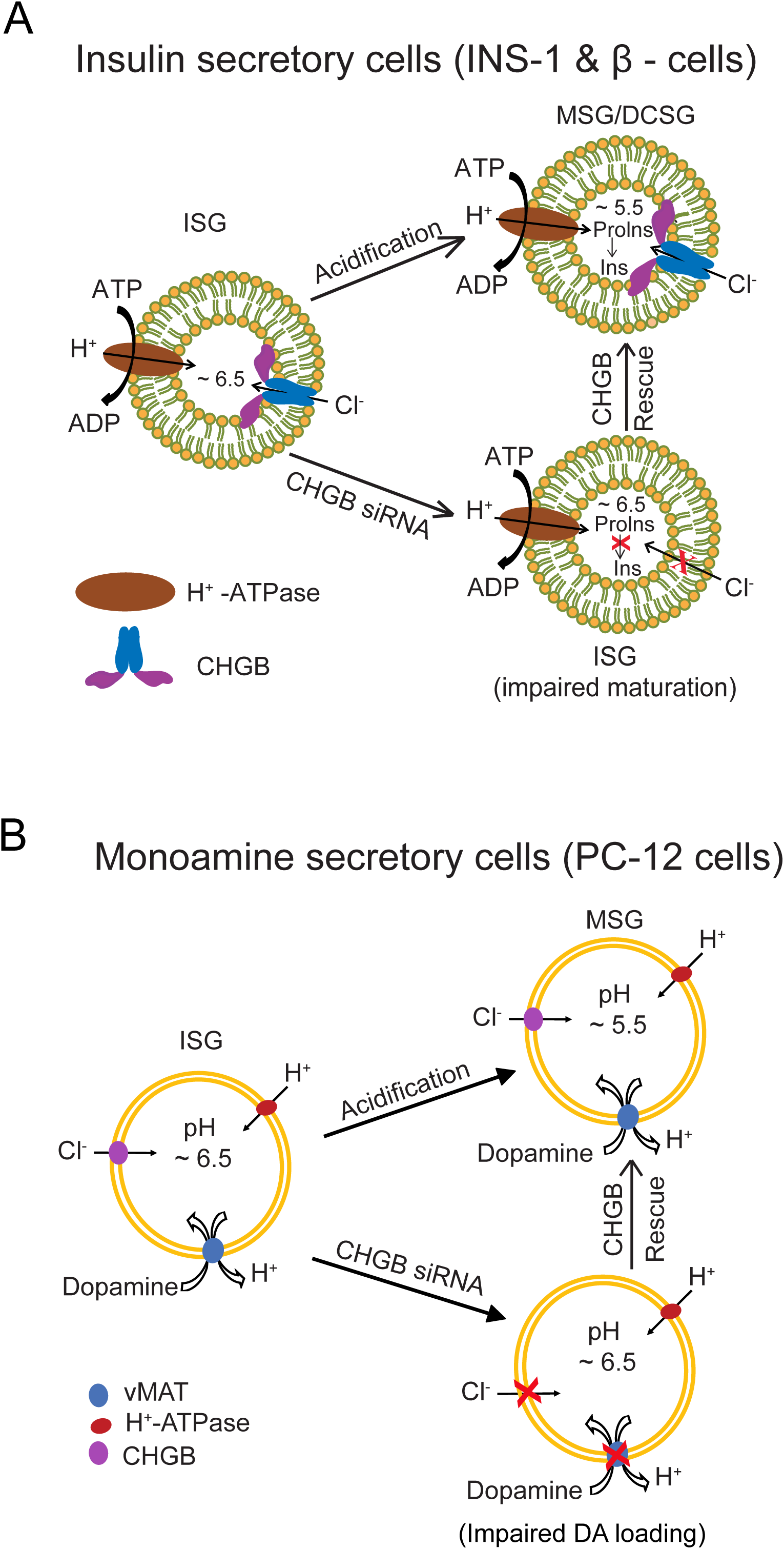
A working model for the CHGB anion channel serving as the long-sought Cl^-^ shunt pathway in regulated secretion. (**A**) CHGB channel, together with vesicular H+-ATPase, supports normal acidification of secretory granules in insulin-secreting cells (both NS-1 and primary β-cells). CHGB knockdown impairs acidification and delays cargo maturation (proinsulin-to-insulin conversion), and overexpressing CHGB restores acidification and insulin maturation. (**B**) In monoamine-secretory cells (PC-12 or dopaminergic neurons), the proton-gradient created during granule acidification is used for H+-coupled loading of catecholamines (e.g. dopamine) into secretory granules. CHGB knockdown impairs acidification and dampens catecholamine loading. Its overexpression restores the monoamine loading.

Consistent with impaired granule acidification in *Chgb*^-/-^ mice, hyperproinsulinemia and insulin secretion defects under glucose challenge were reported before in these animals ^13^. A significant decrease of catecholamine content was observed in chromaffin granules ^50^, and the deficiency was attributed to an unknown intracellular mechanism leading to saturation of catecholamine- binding capacity at the luminal side ^50^. Instead, our model depicts a new mechanism --- the lack of CHGB channel activity de-acidifies chromaffin granules and thereby diminishes the driving force for catecholamine loading and decreases steady-state concentration of monoamines inside the granules (Fig. 7B). Furthermore, under the same principle of electroneutrality, the CHGB Cl^-^ conductance, which conducts well in both directions (Fig 2B), may facilitate Ca^2+^ release from secretory granules or Ca^2+^ influx from the extracellular side, both of which could accelerate RRG release because elevation of local [Ca^2+^] is a prerequisite for granule exocytosis (Figs. 2A, 2C). Our simplified mechanism cannot explain why only a small fraction (0.5% in mouse β-cells) of mature DCSGs are readily releasable ^9^. Moreover, in *Chgb*^-/-^ mice, compensatory expression of other conductances or time-integral of slow granule maturation (due to membrane leaks) may partially mitigate the deacidification phenotypes, which makes *Chgb*^-/-^ mice seemingly healthy under the breeding conditions. Given the importance of regulated secretion, existence of intrinsic compensation is not surprising. Identification of these compensatory molecules and mechanisms will be interesting for future studies.

Our observations in cultured neuroendocrine cells and primary β-cells explain the main phenotypes of *Chgb*^-/-^ mice --- hyperproinsulinemia and decreased catecholamine content in chromaffin granules ^12,13,50^. Future studies will need to examine whether the missense CHGB mutations genetically associated with Type 2 diabetes, neuroendocrine or neurodegenerative diseases alter the channel function, granule acidification and maturation, or release ^12^. Overexpression of CHGB and CHGA has been observed in a large array of human neuroendocrine cancer tissues ^72^, but their functional significance remains a mystery. Verification of a causative relation between altered CHGB channel function and these human diseases will likely make the channel a good target for new therapeutics.

### CHGB differs from all other dimorphic proteins in forming highly selective anion channels

Multiple soluble proteins can be inserted into membranes and form large non-selective conductances that are usually pathogenic or cytotoxic. Many hemolysins are pore-forming toxins. *S. aureus α*-toxin, for example, is secreted as monomers and oligomerizes in host cell membranes to form multimeric complexes which contain non-selective, large-conductance, beta-barrel pores ^73,74^. C-type lectins are released from small intestine epithelial cells as monomers and are activated by proteolytic removal of their N-terminal peptides before forming hexameric pores in membranes of *G^+^* bacteria ^75,76^. VopQ, a pathogenic effector protein of the *Vibrio* species, is released inside the infected cells and punctures lysosomal membranes via forming non-selective large-conductance channels to cause cell demise ^77,78^. Other examples include CLIC family proteins (whose physiological functions remain not fully settled) ^79–83^, the membrane-attack complex formed by the C5b/C6-9 in the complement pathways ^84–89^, the channel forming colicins ^90–92^, the pore forming Gasdermin D (GSDMD) in pyroptosis ^93–97^, the pore-forming BAK or BAX in outer mitochondrial membrane that releases cytochrome C during apoptosis ^98–101^, the membrane-permeabilization through the pores formed by phosphorylated MLKL during necroptosis ^102–106^, the pore-forming microbiocidal defensins ^107–109^, etc. All the membrane-spanning pores formed by these well-studied proteins are non-selective and almost always cytotoxic via breakdown of transmembrane ionic gradients and loss of nutrients, ATP and other metabolites. Contrastingly, the CHGB Cl^-^ channels differ from all of them because of high anion selectivity and the physiological, instead of pathological, roles in the regulated secretory pathways in exocrine, endocrine and neuronal cells. The more stringent anion selectivity is obviously important in order to prevent the breakdown of ion gradients across membranes of the regulated secretory pathway, and minimize the post-exocytotic loss of organic anionic metabolites that are abundant inside these cells.

### CHGB’s physiological functions in granule biogenesis, maturation and release

Being one of the most abundant proteins and present almost in every human tissue, CHGB was previously assumed to participate in every step of the regulated secretory pathway. Its N-terminal Cys-loop serves as a sorting signal ^110,111^. Its Ca^2+^-induced aggregates ^14^, together with other proteins and cargos at the TGN, may induce membrane budding for granule biogenesis. Aggregation may keep a major fraction of CHGB in its soluble form (Figs. 1D-E). High- concentrations of Ca^2+^ or Zn^2+^ inside secretory granules may induce CHGB aggregation in both membrane-bound and soluble states (Fig. 3).

The CHGB anion channel serves an important role in normal granule maturation. Without its N- terminal sorting signal, the rest of CHGB might still form a channel, but would be mis-delivered. CHGB likely uses a strong sorting signal for proper delivery of the channel-forming portion. The CLC3 Cl^-^/H^+^ exchanger, very probably being absent from insulin-secretory granules ^112^ and contributing little anion flux due to its tiny conductance and strict outward rectification [comparing Fig. 2B with the data by ^113,114^], functions differently in other intracellular compartments. However, even without residing in membranes of secretory granules, CLC-3 in cell surface or in endosomal membranes might still affect proton-pumping or chloride flux via physical interactions and facilitate Ca^2+^ influx via charge balance, which could explain its permissive role for granule release as observed in *Clc3-null* mice ^115,116^. Because CHGB and CLC-3 are important for granule maturation and granule release, respectively, both *Clc3*^-/-^ and *Chgb*^-/-^ mice exhibited hyperproinsulinemia and lower catecholamine content in secretory granules ^12,13,50,115,116^. In *Clc3^-/-^* β-cells, secretory granules, being retained in the cytosol for a much longer time, might fuse with lysosomes or endo-lysosomes that could not be properly acidified without CLC-3 and become partially deacidified ^7,115,117–120^. Further experiments could be performed to test this LRO-based mechanism.

CHGB may directly interact with IP_3_Rs and regulate Ca^2+^ release from the granules and in turn calcium-dependent exocytosis ^35,121,122^. The soluble CHGB may interact with IP_3_Rs from granular interior and regulate Ca^2+^ release ^123,124^. Further studies are needed to understand the proposed gating effects of CHGB on IP_3_Rs before granule exocytosis.

With its single channel conductance orders of magnitude higher than that of CLC-3 or -5, CHGB channels retained on cell surface become the dominating Cl^-^ conductance during endocytosis of granular puncta (Fig. 1A) ^42^. Recent high-resolution structures of CLC channels made it quite certain that CLC-3 is a transporter, not a channel ^115,125–145^. CLC-3 was observed in early and late endosomes and synaptic vesicles ^113^ whereas CLC-5 was detected in endocytic vesicles, especially in kidney proximal tubules ^146^. After CHGB on the cell surface is endocytosed, its anion channel function would co-exist with CLC-3 or CLC-5 in endosomes and may be suppressed inside the endocytic compartments. There is hence a site where the secretory granules and the endosomes may interact with each other. Whether CHGB functions as anion channels in the endosomes or during its recycling to *Golgi* membranes, LROs or secretory granules remains an open question.

### Relations between CHGB and other proposed granular ion channels

Besides IP_3_Rs, there are at least three distinct views on ion channels in secretory granules. First, a large-conductance Ca^2+^-activated K^+^-channel (BK), a few other K^+^ channels and a 250 pS Cl^-^ channel (450/150 mM Cl^-^) were reported in isolated chromaffin granules ^34^. Second, no significant permeability of K^+^, Na^+^, Mg^2+^ or Ca^2+^ in chromaffin granules was observed four decades ago ^26,27^. These two views are at odds with each other, but could be reconciled in consideration of low detection sensitivity of the flux assays performed in the earlier study ^27^ or possible contamination of the isolated chromaffin granules used in the more recent work ^34^. Our whole-cell recordings from PC-12 cells (Fig. 2A) detected mainly anion currents, suggesting that in secretory granules the anion channels are probably much more abundant than the proposed K^+^ channels, if any. The 250 pS Cl^-^ channel might represent the CHGB channel as its conductance could double from ∼125 pS when [KCl] is increased from 150 to 450 mM, which awaits verification using specific inhibitors or genetic manipulation.

Third, Kir6.1, ATP-sensitive K^+^/Cl^-^ channels and CLC-1/2 channels were reported from zymogen granules, but the recorded Cl^-^ channels showed properties completely different from the same channels recorded from cell surfaces ^37,38,114^. These conflicting results might have stemmed from experimental difficulties in achieving high purity of zymogen granules with no contamination from other cellular membranes. In contrast, our genetic manipulations had higher specificity and thus avoided technical limitations of previous studies. We detected anion channels from released granules (Fig. 2) and probed specifically the physiological roles of native CHGB channels in granule maturation and acidification (Figs. 4-6). Our findings set a stage for future studies of the proposed relations between CHGB and other ion channels in regulated secretion.

## CONCLUSIONS

Our data from six different angles collectively support the main conclusion that native CHGB proteins in bovine, rat and mouse neuroendocrine cells all form a chloride channel inside secretory granules and on cell membranes, and this channel function is necessary for normal granule acidification and cargo maturation in neuroendocrine cells. The well-conserved CHGB proteins from zebra fish to human constitute a new family of intracellular anion channels that serve the Cl^-^ shunt pathway proposed more than four decades ago and are essential to normal acidification of intracellular compartments related to regulated secretion and probably the recycling of the granular components in various types of CHGB-expressing cells. Potential CHGB functions in ER and Golgi apparatus, budding of nascent granules and homotypic fusion of ISGs as well as the resorting of ISG contents remain open for future studies.

## METHODS: (Details available in Supplemental Information)

1. Molecular cloning of CHGB and its different mutants in pFastbac1
2. Knockdown of CHGB in neuroendocrine cells and rescue by overexpressing CHGB or CHGBΔMIF
3. Detection of insulin and proinsulin in INS-1 cells
4. Western blotting of vesicular ATPase subunit A2 (ATP6V0A2)
5. ELISA assay for detecting dopamine released from PC-12 cells
6. pH measurements in intracellular acidic compartments using DND-160
7. Isolation of pancreatic islets from wild-type and CHGB-knockout mice
8. Purification of native bCHGB from bovine pancreas
9. Electrophysiological recordings from PC12 cells before and after depolarization-induced Ca^2+^-dependent release of secretory granules.
10. High-pressure freezing and immuno-Electron Microscopy (HPF immuno-EM)
11. Quantitative analysis of HPF immuno-EM data
12. Preparation of monolayer cultures of pancreatic beta-cells.
13. *In situ* calibration of ratiometric pH measurement using NPY-ClopHensor
14. Fluorescence imaging and analysis using NPY-ClopHensor
15. Granule biogenesis assay of CHGBΔMIF in CHO cells

## DECLARATIONS

### Ethics approval and consent to participate

The authors confirm that all experiments with mice were performed by following protocols approved by the IACUC committee at UF.

### Consent for publication

All authors agree to publish this paper.

### Availability of data and materials

All raw data are available upon request. We will share the constructs and the information of the animal model with those who will make requests.

### Author Contributions

Q.-X.J. designed and oversaw the experimental studies, analyzed the results with all co-authors. G.Y. designed and performed all the molecular biology, biochemistry and cell-based experiments and analyzed acquired data. H.W., Q.W. and M.X.Z. conducted the electrophysiological experiments in PC-12 cells and analyzed data together with Q.-X.J. M.A. and G.Y. performed the dissection of mice and isolation of pancreatic islets with C. M’s advice. W.H and E.A.P provided advice and assistance in preparation of monolayers of pancreatic beta-cells. J.O., S.C. and P.V. did HPF-immuno-EM and data analysis. All authors contributed to data analysis and manuscript writing.

## Funding and acknowledgements

Details are in the supplementary information.

